# Assessing fecal metaproteomics workflow and small protein recovery using DDA and DIA PASEF mass spectrometry

**DOI:** 10.1101/2024.03.13.584844

**Authors:** Angela Wang, Emily E F Fekete, Marybeth Creskey, Kai Cheng, Zhibin Ning, Annabelle Pfeifle, Xuguang Li, Daniel Figeys, Xu Zhang

**Affiliations:** Regulatory Research Division, Biologic and Radiopharmaceutical Drugs Directorate, Health Products and Food Branch, Health Canada, Ottawa, Canada; School of Pharmaceutical Sciences, Faculty of Medicine, University of Ottawa, Ottawa, Canada; Department of Biochemistry, Microbiology and Immunology, Faculty of Medicine, University of Ottawa, Ottawa, Canada

**Keywords:** Fecal metaproteomics, microbiome, mass spectrometry, differential centrifugation

## Abstract

**Aim:** This study aims to evaluate the impact of experimental workflow on fecal metaproteomic observations, including the recovery of small and antimicrobial proteins often overlooked in metaproteomic studies. The overarching goal is to provide guidance for optimized metaproteomic experimental design, considering the emerging significance of the gut microbiome in human health, disease, and therapeutic interventions.

**Methods:** Mouse feces were utilized as the experimental model. Fecal sample pre-processing methods (differential centrifugation and non-differential centrifugation), protein digestion techniques (in-solution and filter-aided), data acquisition modes (data-dependent and data-independent, or DDA and DIA) when combined with parallel accumulation-serial fragmentation (PASEF), and different bioinformatic workflows were assessed.

**Results:** We showed that, in DIA-PASEF metaproteomics, the library-free search using protein sequence database generated from DDA-PASEF data achieved better identifications than using the generated spectral library. Compared to DDA, DIA-PASEF identified more microbial peptides, quantified more proteins with less missing values, and recovered more small antimicrobial proteins. We didn’t observe obvious impacts of protein digestion methods on both taxonomic and functional profiles. However, differential centrifugation decreased the recovery of small and antimicrobial proteins, biased the taxonomic observation with a marked over-estimation of *Muribaculum* species, and altered the measured functional compositions of metaproteome.

**Conclusion:** This study underscores the critical impact of experimental choices on metaproteomic outcomes and sheds light on the potential biases introduced at different stages of the workflow. The comprehensive methodological comparisons serve as a valuable guide for researchers aiming to enhance the accuracy and completeness of metaproteomic analyses.

## INTRODUCTION

The human gut microbiome contains an estimated 100 trillion microorganisms, including bacteria, fungi, protozoa, and viruses, which interact with each other and their host to foster a complex and dynamic environment ^1-3^. The symbiotic host-microbial relationship of the gut microbiome is crucial to human health and contributes to many biological processes, such as metabolism, immunomodulation, etc ^4^. Many studies also suggest that microbiome dysbiosis is correlated with, and may lead to the development of neurodegenerative, cardiovascular, metabolic, and gastrointestinal diseases among others ^3,5-7^. With the emerging importance of the gut microbiome in human health, disease, and therapeutics, studies on the microbiome, its taxa, and its products have become increasingly significant ^8^.

Given the extremely high complexity of the microbiome, meta-omics approaches including metagenomics, metatranscriptomics, metabolomics, and metaproteomics, are commonly used in studying the microbiome composition and functions ^9,10^. Among the different –omics approaches, metaproteomics uses a mass spectrometer to directly measure the protein expressions as well as post-translational modifications (PTMs) of the microbial community ^11^. Mass spectrometry (MS) analysis can be conducted with a data-dependent acquisition (DDA) or data-independent acquisition (DIA) strategy. DDA-based metaproteomics is commonly used due to its easy set up and analysis, flexibility, breadth of detection, and ability to relatively quantify chemically labelled peptides ^12^. In a DDA mode, the most abundant ions from MS 1 scan will be selected and fragmented during tandem MS scans, however, this data acquisition mode can risk losing information on the other less abundant peptides particularly in complex samples such as microbiomes, which limits the depth, sensitivity, and reproducibility of metaproteomic data ^13,14^. Contrastingly, DIA can sample all the peptides within the selected mass range and therefore theoretically can detect lower-abundance peptides in complex samples ^15^. In the past few years, the application of DIA-MS based proteomics was profoundly expanded due to these advantages, the advancement of bioinformatics tools, such as DIA-NN, and advanced instrumental developments, such as DIA-PASEF and Astral MS analyzer ^16,17^. More recently, the application of DIA-MS in metaproteomics has been reported, demonstrating great potential in increasing the depth of identification and accuracy of quantifications ^18^.

One advantage of metaproteomics is the capability to measure non-bacterial components, including proteins originating from the host as well as viral, fungal and archaeal species without the need of additional experimental efforts ^19,20^. This is particularly ideal for trans-kingdom, host-microbiome interaction studies ^21,22^. The intestinal lumen is the home of diverse biotic and abiotic components, including host secreted proteins such as antimicrobial proteins as well as proteins produced by microbes themselves, such as small microbial proteins. These small proteins/polypeptides, including a high proportion of antimicrobial peptides, are implicated in cell-signaling, transport, enzymatic activities, antitoxin systems, pathogenic colonization resistance, etc ^23-25^. Theoretically, these small proteins present in the intestinal samples can be identified in metaproteomics data, however, in practice, their identification is often overlooked due to suboptimal sample preparation and bioinformatics annotation steps which loses these portions of protein components^26,27^. For example, the secreted small antimicrobial peptides from either the host or microbes may be lost during differential centrifugation, a commonly used sample pre-processing step to purify fecal samples prior to protein extraction.

It is well recognized that different sample processing methods such as the use of differential centrifugation, protein extraction methods, protein digestion methods, as well as differences in data analysis workflows can yield different results and metaproteomics insights ^18,20,27-29^. We have previously reported that physical disruptions using bead beating or ultrasonication are needed to optimally extract proteins from Bacillota (previously Firmicutes) ^29^. Tanca et al. showed that stool pretreatment by differential centrifugation significantly impacted metaproteomic observations ^28^. Yet there is still a need of further evaluation of the bias introduced by different steps of the experimental workflow as well as the emerging DIA data acquisition mode; and how they can contribute to the recovery of otherwise overlooked components, such as small antimicrobial proteins. Therefore, in this study, we conducted a comprehensive comparison of fecal sample preparation, protein digestion, data acquisition mode and bioinformatic workflow using mouse feces to serve as a guide for metaproteomic experimental design.

## RESULTS AND DISCUSSION

### Evaluating bioinformatics workflows for DIA-PASEF metaproteomics

To evaluate fecal sample preparation workflows, a pooled fecal sample from C3H/HeN female mice was crushed into a powder, homogenized and aliquoted for either differential centrifugation (DC) or direct protein extraction with non-differential centrifugation (NC) workflow (Figure 1A). To assess the potential impacts of protein digestion methods, both DC and NC protein lysates were subjected to in-solution trypsin digestion (following acetone precipitation for detergent removal), filter aided sample preparation (FASP) with 10kDa molecular weight cutoff filter (FASP10), or 3kDa filter (FASP3). All comparisons were conducted with 5 replicates with a total of 30 peptide samples for MS analysis on a timsTOF Pro 2 mass spectrometry system using both DDA- and DIA-PASEF acquisition modes (Figure 1A).

**Figure 1.**
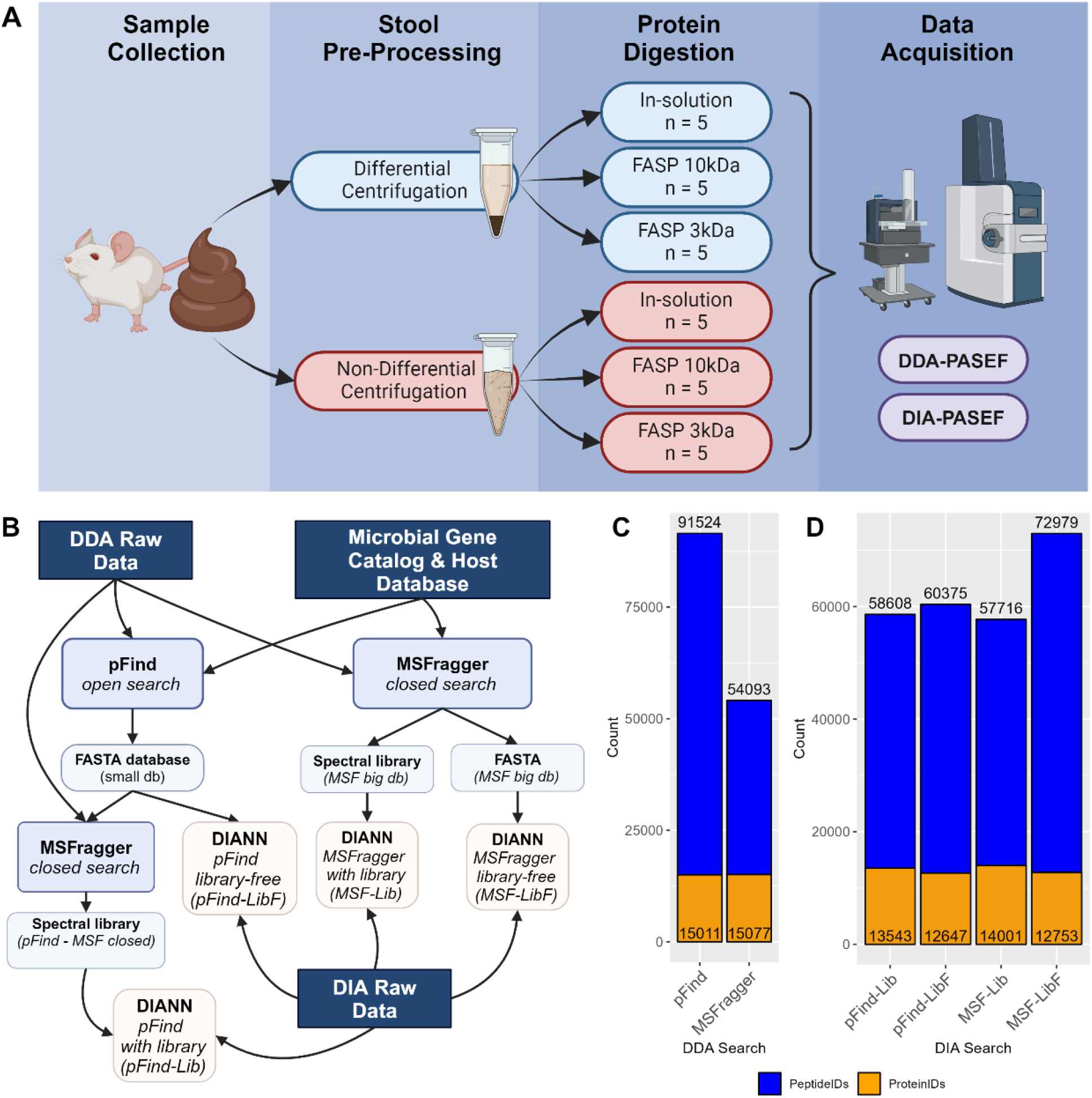
Experimental Overview and evaluation of bioinformatics workflow. Flowchart depicting mouse fecal sample processing workflow and MS data acquisition (A). Flowchart depicting different bioinformatic workflows tested in this study (B). Histograms of the total number of peptides and proteins identified in each bioinformatics data processing workflow (C).

A major challenge for DIA metaproteomic data analysis is that bioinformatics tools, such as DIA-NN, cannot handle large databases. Currently, most workflows utilize DDA data of the same or representative samples to generate a spectral library from the original protein databases. We therefore first evaluated the use of two widely used database search engines for DDA data, pFind (open search) and MSFragger (closed search with split databases), to generate reduced databases from the mouse gut Microbial Gene Catalog database (Figure 1B). The pFind open search identified 91,524 peptides corresponding to 15,011 protein groups in total, while MSFragger identified 54,093 peptides corresponding to 15,077 protein groups (Figure 1C). DIA-NN searches with spectral library or library free modes from either pFind or MSFragger reduced databases were then conducted to see which pipeline for DIA data processing yields the best identification results. Briefly, protein sequences were extracted from both search outputs to generate reduced FASTA databases for DIA-NN search with library-free mode (pFind-LibF for using pFind generated protein database, and MSF-LibF for using MSFragger generated database). Spectral libraries were generated by MSFragger with either full database or pFind-generated reduced protein database, and were used for DIA data analysis (MSF-Lib and pFind-Lib, respectively). As shown in Figure 1D, all DIA-NN search inputs yielded similar protein group numbers, falling between 12,647 and14,001. The main difference in identification was found in the peptide level, where all workflows tested resulted in approximately 58,000-60,000 peptides except for MSF-LibF which identified 72,979 peptides in total for the DIA-PASEF dataset. Based on these evaluations, we then chose MSF-LibF workflow for DIA-PASEF data processing, and to be consistent the MSFragger LFQ-MBR quantification using the full database (with 10 splits) was used for the analysis of DDA-PASEF dataset.

### DIA-PASEF metaproteomics achieved better identification and quantification

We first compared the DDA-PASEF and DIA-PASEF data acquisition modes in terms of peptide and protein identification as well as protein quantification in fecal metaproteomics. Across all fecal sample preparation methods, there was a consistent increase in peptides and protein groups identified when samples were run using DIA-PASEF mass spectrometry compared to those run with DDA-PASEF mode (Figure 2A-B). DC and NC preparation methods showed comparable identification of both peptides and protein groups in this study, however, the digestion method using a FASP-10kDa approach had a slight edge over other digestion methods tested in DC preparations, and in-solution digestion had a slight edge in NC samples (Figure 2A-B). DIA-PASEF runs for DC samples achieved the highest number of precursor identifications with nearly 50000 precursors per sample (identifications ranging from 36912 to 49928 precursors, 35324 to 46665 peptides, 9598 to 10705 protein groups), while nearly 21000 peptides (identifications ranging from 14029 to 20852 peptides, to 5068 to 6891 protein groups) were identified for the same samples with DDA-PASEF runs. DIA-PASEF runs for NC samples achieved a competitive number of precursor identifications with nearly 45000 precursors per sample (identifications ranging from 17681 to 44915 precursors, 17457 to 42646 peptides, 7002 to 11031 protein groups), while up to almost 20000 peptides (identifications ranging from 6334 to 19389 peptides, 3292 to 7492 protein groups) were identified for the same samples with DDA-PASEF runs. The filter-assisted sample preparation when using the FASP-3kDa columns showed comparable identification rates when used on DC samples compared to other digestion methods, but led to a significant drop in peptide and protein group identifications when used with NC samples (Figure 2A-B).

**Figure 2.**
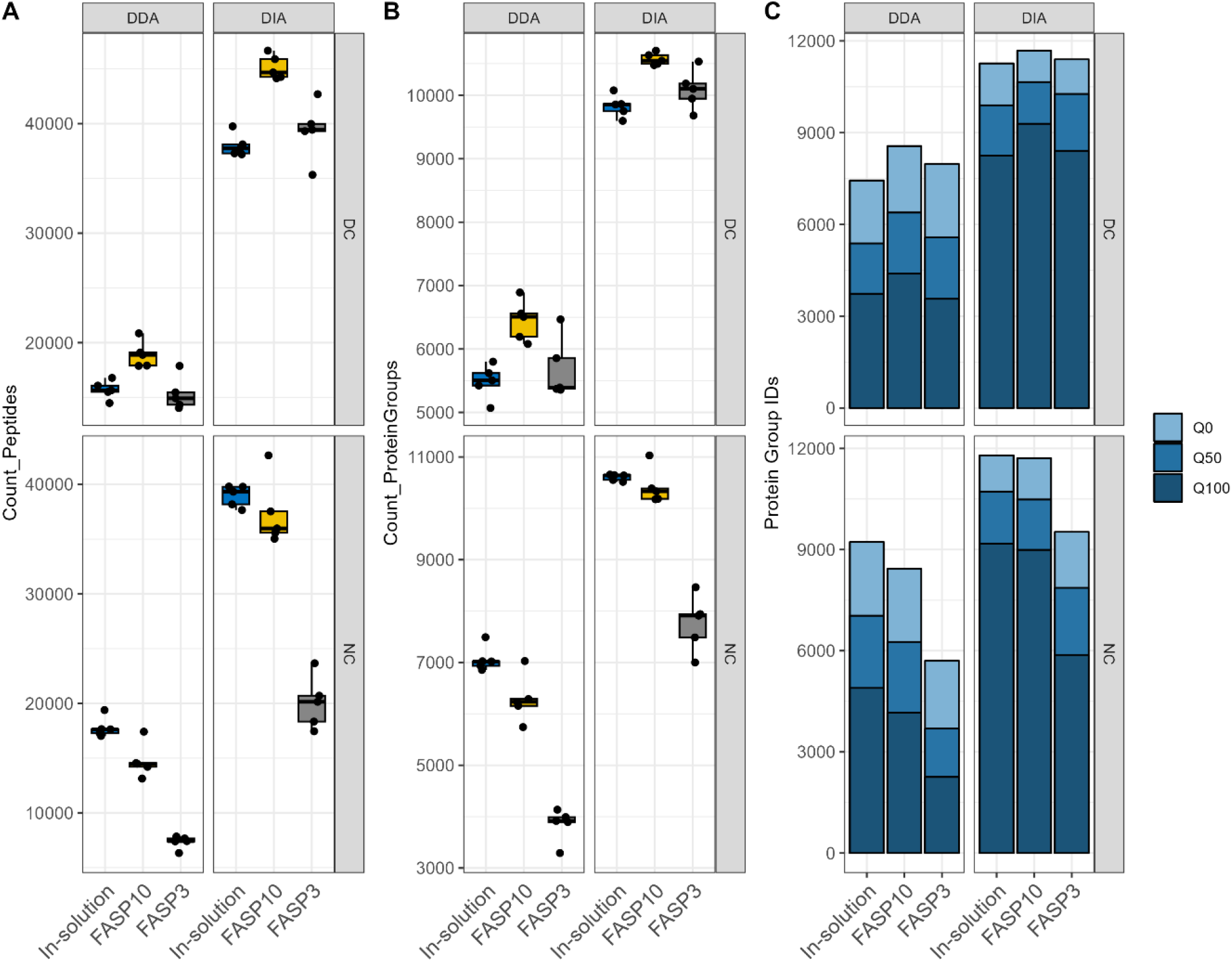
Comparison of identification and quantification abilities of DDA and DIA-PASEF approaches. Number of peptides identified by DDA and DIA-PASEF (A) mass spectrometry methods. Number of protein groups identified by DDA and DIA-PASEF (B) mass spectrometry methods. Number of quantitated protein groups in any, half, or all (Q0, Q50, Q100 respectively) samples in each group identified by DDA- or DIA-PASEF (C) mass spectrometry.

Impacts of MS data acquisition and sample preparation on quantification were then assessed with the amount of missing values across samples in each group. In Figure 2C, Q0 indicates a protein group with quantified intensity in at least 1/5 replicates in the group, Q50 is present in at least 3/5 replicates, and Q100 is present in all replicates. Along with an increased number of protein group identifications in DIA-PASEF data across all sample preparation methods, there is a distinctly higher proportion of protein groups quantified in all sample replicates (Q100) for DIA-PASEF data when compared with DDA-PASEF, in particular for samples prepared with in-solution digestion and FASP-10K column (73% - 80% in DIA vs. 49% - 53% in DDA; Figure 2C). Therefore, DIA-PASEF mass spectrometry methods for mouse metaproteomic samples from various preparations provide more identifications with fewer missing values, providing better quantitative analysis potential.

### DIA-PASEF for non-enriched samples yielded highest recovery of small proteins and antimicrobial peptides

When conducting the database search, we included both microbiome and host protein sequences. When looking at host proteins, DIA-PASEF mass spectrometry methods identified a higher number of host proteins than DDA, as expected since DIA-PASEF identified more proteins in total. Despite an increased diversity of host proteins identified in DIA-PASEF dataset, the relative abundance of host proteins did not show a significant difference between both data acquisition and protein digestion methods (Supplementary Figure 1). Differential centrifugation does not appear to have a major impact on the identification count of host proteins or their relative abundances, which might be due to the relatively low amount of host protein (less than 18% in abundance) in the feces of healthy mice that were used in this study (Supplementary Figure 1). The most abundant host proteins identified in the fecal samples include those involved in digestion function and antimicrobial activity, such as regenerating islet-derived protein 3-beta (Reg3β) and immunoglobins, which may bind tightly to the microbe surface and thereby become enriched with microbial cells.

Next, we evaluated whether different sample preparation methods and MS data acquisition modes could impact the identification of small proteins that are commonly overlooked in classical metaproteomics. Among all the identified proteins in this dataset, there were 31 proteins groups with ≤ 50 amino acids and 845 with ≤ 100 amino acids. We selected a 100 amino acids cutoff in this study for the purpose of comparing between groups. There were 200-600 small proteins identified per sample, with DIA-PASEF yielding higher identifications than DDA-PASEF mode (Figure 3A). Differential centrifugation reduced the number of small protein identifications when the extracted proteins were digested using in-solution and FASP-10kDa methods, but not for those with FASP-3kDa method. Considering the sum relative abundance of those small proteins, the type of digestion, FASP column size, and MS acquisition mode all present no to minimal differences (Figure 3B). We then performed functional and taxonomic annotation using GhostKOALA for these identified small proteins, which showed that they are significantly enriched in genetic information processing and lipid metabolism pathway (Figure 3E-F and Supplementary Figure 3). In addition, the identified small proteins in both DDA and DIA datasets were significantly enriched in undefined taxa (adjust P value of 1.59E-68 and 1.08E-45 for DDA and DIA datasets, respectively) from the microbiomes (Supplementary Figure 4). Altogether, this study showed that the use of differential centrifugation depleted the abundance of small proteins within sample sets across both DDA and DIA acquisition modes and protein digestion methods, suggesting that the NC method is superior when targeting and investigating small proteins specifically. It is worth mentioning that NC method also has the advantage of significantly shortened sample preparation step and time, and thereby reduce the sample-to-sample variations introduced during sample preparation.

**Figure 3.**
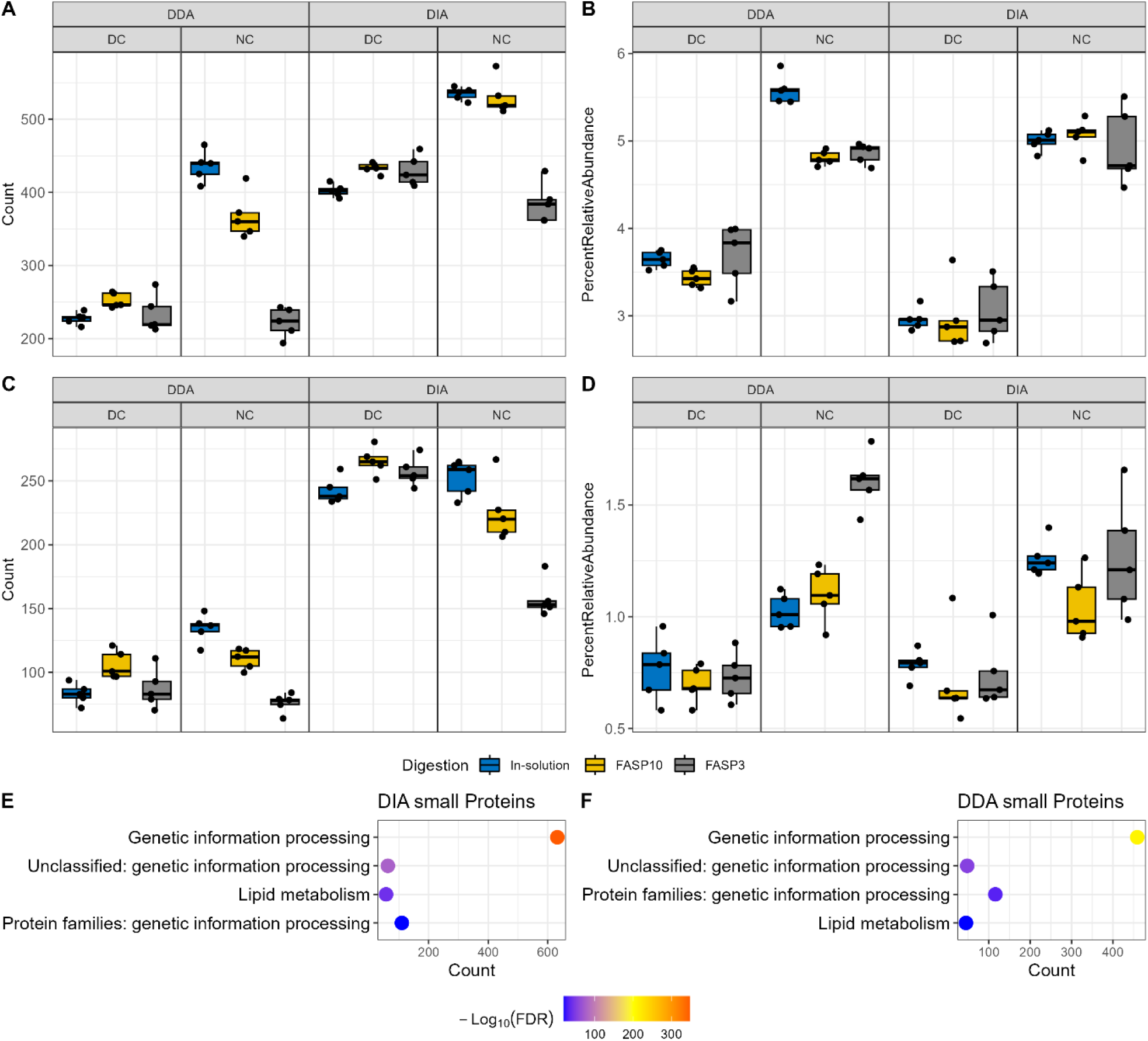
Quantitation of small proteins and antimicrobial peptides in mouse fecal samples. Count (A) and relative abundance (B) of small proteins in samples using a 100 amino acid cutoff. Count (C) and relative abundance (D) of AMPs when searched against the AMPsphere database. Functional enrichment analysis of identified small proteins in DDA-PASEF dataset (E) and DIA-PASEF dataset (F), using all identified proteins as background in DDA and DIA dataset, respectively. Functional annotation was performed using GhostKOALA, and only significantly enriched functions were shown (adjusted P value ≤ 0.05).

**Figure 4.**
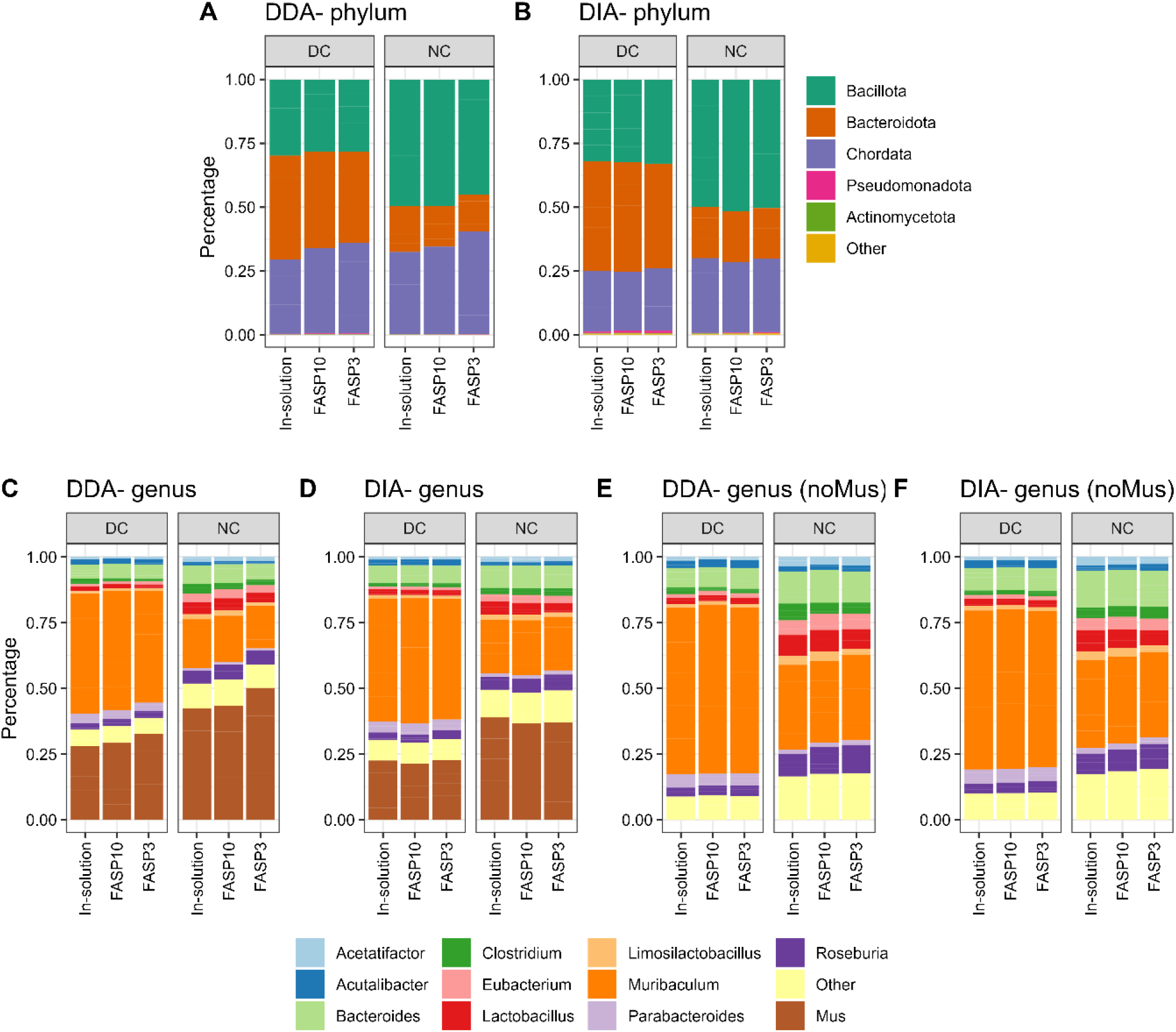
Taxonomic profiles of mouse fecal metaproteome. Phylum level (A-B), genus level with host (Mus; C-D) or without host (E-F) were shown. The group mean relative abundance of phyla or genera was used for plotting. Top 5 phyla or top 10 genera were shown with the remainder grouped as ‘Other’ in the stacked plots.

Previous metagenomics data mining identified small open reading frames (sORF) encoding >4,000 small proteins (≤ 50 amino acids) in the human microbiome with the majority having no known function ^23^. Although the human microbiome may not fully encapsulate the small proteins that would be present in a mouse microbiome, the overlap that exists can help reinforce the evaluation of the impacts of different sample preparation methods and MS data acquisition mode on small protein identification. To enable calculation of the relative abundance and mitigate false discovery, all identified mouse gut microbial peptide sequences in this study were combined with the predicted small protein sequences for database search for both DDA and DIA-PASEF data using MSFragger and DIAN-NN, respectively. A consistent and large increase in small protein identifications in all sample preparation methods was observed when samples were run using a DIA-PASEF mode than with DDA-PASEF regardless of upstream sample preparation workflows (Supplementary Figure 2). NC in combination with FASP-3kDa digestion workflow tends to identify the lowest number of small proteins which may be due to the overall low protein identifications in this group, however the small proteins identified represented the largest percentage in abundance in samples. In fact, for both in-solution and FASP digestion workflows, NC leads to increased abundance of small proteins regardless of protein digestions methods and MS acquisition modes, suggesting that differential centrifugation depletes small proteins within the sample. Altogether, the findings in this study suggest that direct fecal lysis for protein extraction, followed by digestion with either in-solution or FASP-10kDa workflows, and quantified using DIA-PASEF presented the highest number of small protein identifications while maintaining a high relative abundance within the sample.

Small proteins are implicated in diverse functions of the microbiome, including defense against other microbes or pathogens ^30^. Among the predicted small proteins in the human microbiome, around 30% were predicted to be secreted or transmembrane proteins and 39 protein families were predicted to be novel antimicrobial peptides (AMPs) ^23^. By using a deep learning approach, Ma et al. identified 2349 potential AMPs from human microbiome metagenomic data and preliminary biological validation showed >80% positive rate ^25^. More recently, Santos-Júnior et al. leveraged a vast dataset of more than 1.5 million metagenomes or microbial genomes of diverse origins to establish a prokaryotic AMP database, AMPSphere ^31^, which consists of 863,498 predicted AMPs. To evaluate the impacts of a metaproteomic workflow for AMP identification, we re-searched both DDA and DIA datasets using the AMPSphere database concatenated with all identified gut microbial peptide sequences. Consistent with total and small protein identification, DIA-PASEF showed a higher AMP identification (200 ∼ 300 AMPs, excluding the FASP-3K group) when compared to DDA-PASEF methods (50 ∼ 150 AMPs) (Figure 3C). When looking at the relative abundance of AMPs within each sample, it appears as though differential centrifugation depletes AMPs (Figure 3D). The findings suggest that NC samples with digestion methods of either in-solution and FASP-10kDa followed by MS analysis with DIA-PASEF mode resulted in high AMP diversity observed at a high relative abundance.

### Over-representation of *Muribaculum* in mouse fecal metaproteome with differential centrifugation

We next evaluated whether different sample preparation methods and MS data acquisition mode impact the taxonomic profiles using metaproteomics. By using a threshold of a minimum of 3 distinctive peptides for confident taxon identification, this study identified 12 phyla, 13 classes, 21 orders, 23 families, 62 genera and 96 species in the DDA-PASEF dataset, and 17 phyla, 22 classes, 19 orders, 32 families, 70 genera and 96 species for the DIA-PASEF dataset. Here we selected phylum and genus level for the evaluation. No obvious difference on the abundance distribution of abundant taxa was observed between DDA and DIA datasets at both phylum and genus level (Figure 4). The major differences were observed between DC and NC, which is in agreement with previous studies of both human and mouse microbiomes ^12,28,32^. A higher relative abundance of host proteins was observed in NC groups when compared to DC groups, in particular at the genus level (Figure 4A-B). Interestingly, we observed a difference of host protein percentage between different protein digestion methods in DDA-PASEF data, but not in DIA-PASEF data. On the contrary, the microbial compositions were highly consistent in both DDA-PASEF and DIA-PASEF datasets.

Metaproteomics analysis demonstrated that *Bacillota* (previous *Firmicutes*) and *Bacteroidota* (previous *Bacteroidetes*) were the two predominant phyla in mouse feces, however the relative abundances of these two phyla were dramatically altered by the use of differential centrifugation during sample pre-processing. Marked higher relative abundances of *Bacteroidota* and lower abundances of *Bacillota* were observed in DC samples when compared to NC in both DDA and DIA datasets. This is in agreement with previous studies on mouse metaproteomes ^12,32^. However, the direct opposite of changes was reported in human microbiomes where lower levels of *Bacteroidota* and higher levels of *Baccilota* were obtained when samples were prepared using differential centrifugation ^28^. This difference might be due to the known marked different bacterial genus/species compositions of mice and human microbiomes. As shown in Figure 4E-F, *Muribaculum* was the genus mainly driving the elevation of *Bacteroidota* in DC samples, which represent ∼70% of the microbial abundance in the samples. *Muribaculum* species are known to be dominant in mouse gut microbiota, but were only recently well characterized and demonstrated to have very high host preference (prevalence of 67% in mice compared to 7% in human gut) ^33^. In contrast, in human gut microbiota, the genus *Bacteroides* is usually the most abundant in *Bacteroidota* ^34^. In this study, we found that the relative abundance of *Bacteroides* was lower in DC compared to NC, which is in agreement with the observations in human gut microbiota. It is unknown why *Muribaculum* displayed different responses to DC when compared to their neighboring genera within the sample phylum *Bacteroidota*, but this observation suggests that sample preparation methods need to be optimized for microbiomes of different origins and the microbial species of interest for a particular study.

### Differential centrifugation altered the functional profiles of fecal metaproteomes

Lastly, we evaluated the influence of sample preparation methods and MS acquisition on the functional profiles obtained using fecal metaproteomics. We identified 24 out of the 26 Clusters of Orthologous Gene (COG) categories for both DDA and DIA datasets with 1356 and 1261 COGs, respectively, in this study. As shown in Figure 5, a consistent COG category level composition of fecal metaproteome was observed regardless of protein digestion method and MS acquisition mode. However, both DDA-PASEF and DIA-PASEF datasets demonstrated that the differential centrifugation dramatically altered the observed functional profiles in metaproteomics. These include the marked decrease of functional category J (translation, ribosomal structure and biogenesis) and N (cell motility) by differential centrifugation, while increased category E (amino acid transport and metabolism), M (cell wall/membrane/envelope biogenesis), R (general function prediction only), and P (inorganic ion transport and metabolism). The decrease of functions related to translation and cell motility suggests that the differential centrifugation may result in an under-estimation of microbial species with more active cell proliferation and higher motility. The functional profiles analysis again demonstrated that sample preparation methods need to be optimized for studies with specific objectives or functional pathways of interest.

**Figure 5.**
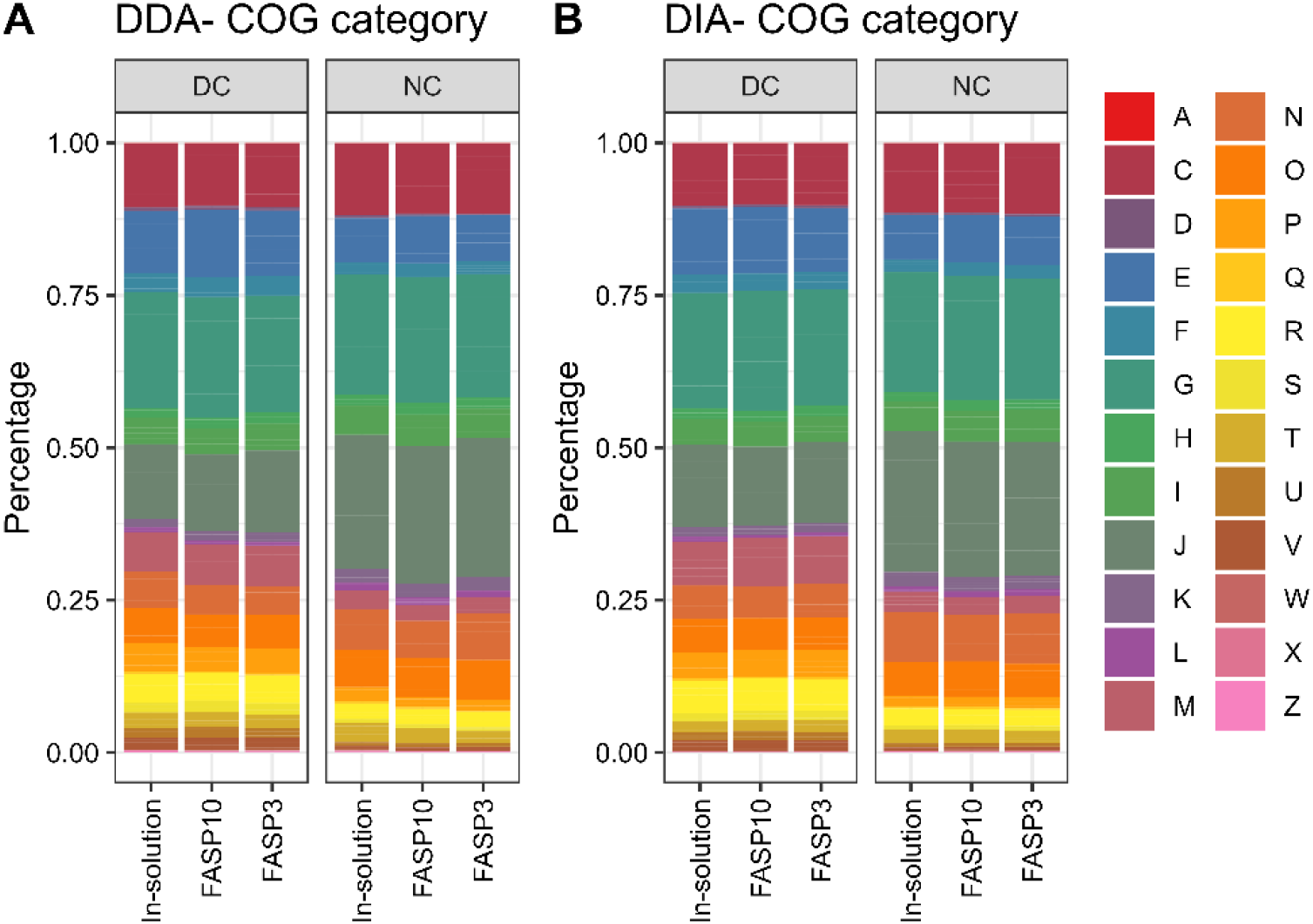
Composition of identified COG categories in mouse fecal metaproteome. The group mean relative abundance of COG category was plotted for DDA (A) and DIA dataset (B), respectively. Each letter represents a COG category according to the Database of Clusters of Orthologous Genes (COGs) https://www.ncbi.nlm.nih.gov/research/cog/.

## CONCLUSION

In conclusion, this study offered a comprehensive comparison of different experimental parameters, including fecal sample pre-processing methods (differential centrifugation and non-differential centrifugation), protein digestion techniques (in-solution and FASP with different molecular cutoff sizes), data acquisition modes (DDA- and DIA-PASEF), and different bioinformatic workflows. Our findings showed that DIA acquisition provided a clear advantage when compared to DDA for identification and quantification of proteins, including small and antimicrobial proteins/peptides, in microbiome samples. We also demonstrated that non-differential centrifugation methods improved the recovery of small proteins and AMPs, and that FASP workflow using 10kDa molecular cutoff filter achieved similar data outputs compared to in-solution digestion, both of which are commonly used in proteomic and metaproteomic studies.

We have previously reported that lysis buffer and protein extraction protocols had major impacts on metaproteomics observations, and the protein extraction protocols with strong detergent sodium dodecyl sulfate (SDS) and ultrasonication achieved the best protein yields and peptide/protein identifications ^29^. Together with this previous study, we highlighted the critical impact of experimental choices on metaproteomic outcomes and shed light on the potential biases introduced at every step of the workflow. The outcomes of this study provide valuable information in standardizing the metaproteomics workflow for applications, such as clinical study, drug development and regulatory assessment, especially for microbiome-based medicinal products.

## METHODS

### Mouse feces collection and differential centrifugation

Mouse fecal samples were collected from nine female C3H/HeN mice purchased from Charles River Laboratories, Senneville, Quebec, Canada. The animal procedures were approved by the Animal Care Committee in Health Canada and performed in accordance with institutional guidelines. Fecal samples were combined and crushed into a homogenous powder. This stock fecal sample was used for both differential centrifugation and non-differential centrifugation workflows.

Differential centrifugation of feces was performed according to previous study ^29^. Briefly, 0.5mL glass beads and 7.5mL cold PBS per gram of fecal sample were added to the samples followed by vortexing and centrifugation at 300g at 4°C for 5 minutes to collect the supernatant. The remaining fecal pellets were extracted three more times with 7.5 mL, 5mL and 5mL cold PBS, respectively, and discard the fecal pellet at the end of the 4 extractions. Once pooled, additional debris from the extractions was removed by three centrifugations at 300g at 4°C for 5 minutes. The supernatant extract was then spun down at 14000g at 4°C for 20 minutes to collect the microbial pellet. The microbial pellet was washed twice with cold PBS by resuspending and centrifuging at 14000g at 4°C for 20 minutes, then frozen until use.

### Protein extraction, trypsin digestion and desalting

#### Sample lysis

Samples were lysed by resuspending 20mg of frozen non-differential centrifuged (NC) fecal pellet in 1mL lysis buffer or ∼120mg of frozen microbial pellet following differential centrifugation (DC) in 1.2mL of lysis buffer. The same lysis buffer was used for all samples, consisting of 4% (w/v) SDS, 8M urea in 50mM Tris-HCl. Sample lysates were sonicated using a QSonica Q700 water-chilled cup-horn sonicator at 50% amplitude, 10 seconds pulse on/off cycle, for 20 min active sonication time at 8°C. The lysate was then centrifuged at 16,000g for 10min at 8°C to remove any non-lysed debris. Protein-containing supernatant was transferred to a new tube and protein concentrations was determined using the Pierce BCA Protein Assay Kit (Thermo Fisher Scientific, Cat #23225) following the manufacturer protocol.

#### In-solution trypsin digestion

For in-solution trypsin digestion, proteins underwent an acetone precipitation by adding 5 volumes of protein precipitation buffer (50% acetone, 50% acetonitrile, 0.1% acetic acid), mixing by inversion and incubated at –20°C overnight. Samples were spun down at 16000g at 4°C for 25 min and supernatant were discarded. The remaining pellet was washed with ice-cold acetone, sonicated with a QSonica Q700 water-chilled cup-horn sonicator at 50% amplitude for 10 seconds, spun down at 16 000g at 4°C for 10 minutes for a total of 3 washes. After brief air dry, the protein pellets were resuspended in 6M urea in 50mM ABC buffer for trypsin digestion. Protein concentrations was determined using the Pierce BCA Protein Assay Kit (Thermo Fisher Scientific, Cat #23225), and 100ug of protein lysates of each sample were reduced with 10mM 1,4-dithiothreitol (DTT) incubated at 850rpm at 56°C for 30 min and alkylated with 20mM iodoacetamide (IAA) for 40 min at room temperature protected from light. Samples were diluted in 50mM ammonium bicarbonate (ABC) buffer to a final urea concentration of 0.6M and then digested with 4ug MS-grade Trypsin / 100ug of protein incubated at 850rpm at 37°C for 20 hours (overnight). The reaction was stopped by adding formic acid to acidify the samples to pH2-3 prior to desalting as described below.

#### Filter-Aided Sample Preparation (FASP) digestion

For filter assisted digestion methods, two different molecular weight cut-off filters were used, namely FASP-10kDa and FASP-3kDa, in this study. For each sample, 100ug of protein lysate was directly diluted to 200uL of 8M urea in 100mM Tris-HCl buffer and added to the pre-rinsed FASP columns. SDS was diluted twice using 200uL of 8M urea in 100mM Tris-HCl buffer each time. Reduction was performed by adding 8M urea in 100mM Tris-HCl buffer containing a final concentration of 20mM DTT and incubated at 850rpm at 37°C for 30 min. After removing the flow-through, samples were then alkylated by 20mM IAA at room temperature in the dark for 30 minutes. To remove excessive IAA, additional 200ul 20mM DTT in 8M urea in 100mM Tris-HCl buffer was added, incubated at room temperature for 2 min, and eluates were discarded. The column was then washed with 8M urea in 100 mM Tris-HCl buffer once and 100mM Tris-HCl buffer for four times prior to adding 200ul 100mM Tris-HCl buffer containing 4ug of trypsin (1ug trypsin : 25ug protein input). The trypsin digestion was performed by shaking at 850 rpm at 37°C overnight. Peptides were eluted by spinning at 16 000 g at room temperature for 20min followed by additional elution with 200ul fresh 100mM Tris-HCl buffer. Both eluents were combined and acidified to a pH of 2-3 using 10% (v/v) formic acid for desalting.

#### Desalting

Desalting was performed using C18 columns (Thermo Scientific, Cat#89870). Columns were activated by adding 100% acetonitrile (ACN), centrifuging at 100g for 1 min for total of 3 times then equilibrated with 0.1% (v/v) FA, centrifuging at 300g for 2 min for 2 times. Samples were loaded to the column and centrifuged at 300g to remove flow-through until all samples were loaded. The desalting column was washed with 0.1% (v/v) FA for 2 times. Desalted peptides were then eluted with 100ul 80% (v/v) ACN/0.1% (v/v) FA buffer for two times by centrifuging at 100g for 1 min for each elution. The eluates containing desalted tryptic peptides were then dried on a centrivap (Labconco, Cat#7810010) and stored at -20°C until LC-MSMS analysis.

### LC-MSMS analysis

Dried tryptic peptides were re-suspend in 0.1% (v/v) formic acid (FA) to a final concentration of 1 ug/ul, and 3ug peptides were loaded for MS analysis using a timsTOF Pro 2 mass spectrometer (Bruker Daltonik, Bremen, Germany) coupled to a nanoElute 2 UPLC system (Bruker Daltonik). The instrument was calibrated prior to analysis with Chip Cube High Mass Reference Standard (Agilent, G1982-85001). A two-column system of HPLC was used consisting of a C8 trap column before separating on a PepSep Twenty-five analytical column (25 cm x 75 μm column packed with 1.9 μm C18 particles) (Bruker Daltonik). Chromatographic separation was achieved at a flow rate of 0.5µl/min over 48 min in linear steps as follows (solvent A was 0.1% formic acid in water, solvent B was 0.1% formic acid in acetonitrile): initial, 2%B; 40min, 35%B; 40.5min, 95%B; 45min, 95%B; 48min 95%B. The eluting peptides were analyzed in either data-dependent acquisition coupled with parallel accumulation serial fragmentation (DDA-PASEF) mode or data-independent acquisition coupled with PASEF (DIA-PASEF) mode in the timsPro 2 mass spectrometer.

For DDA-PASEF mode, a MS survey scan of 100-1700m/z and ion mobility range of 0.85-1.30Vs/cm^2^was performed in the TimsTOF MS. During the MS/MS scan, 4 PASEF ramps were run with an intensity threshold of 2500, target intensity of 20000 and a maximum precursor charge of 5. The TIMS analyzer was operated in a 100% duty cycle with equal accumulation and ramp times of 100 ms each and a total cycle tile of 0.53 s. The collision energy was ramped linearly as a function of mobility from 59 eV at 1/K0 = 1.6 Vs/cm2 to 20 eV at 1/K0 = 0.6 Vs/cm2.

For DIA-PASEF mode, a MS survey scan of 100-1700m/z and ion mobility range of 0.6-1.60Vs/cm^2^ was performed. The TIMS analyzer was operated in a 100% duty cycle with equal accumulation and ramp times of 100 ms each and a total cycle time estimate 1.8 sec. During DIA-PASEF MS/MS scan, precursors with m/z between 400 and 1200 were defined in 16 scans containing 32 ion mobility steps with an isolation window of 26 Da in each step with 1 Da overlap with neighbouring windows. The collision energy for DIA-PASEF scan was ramped linearly from 59 eV at 1/k0 = 1.3 V·s/cm2 to 20 eV at 1/k0 = 0.85 V·s/cm2.

### Bioinformatic data processing

#### Spectral library and reduced database generation

Mouse gut microbial gene catalog database, containing ∼2.6 million nonredundant protein sequences, was downloaded from GigaScience Database (http://gigadb.org/dataset/100114) ^35^. The reviewed mouse UniProtKB database were downloaded from Uniprot (downloaded on 2024/01/01, 17179 protein entries). These two databases were combined as starting original database for spectral library or reduced FASTA database generation with DDA-PASEF dataset using either pFind ^36^ or MSFragger ^37^. For pFind search, MS raw files were first converted from .d to .mgf files using MSconvert (v3.0.23240). The resulted MGF files were then searched using open search mode against the combined database with the following parameters; enzyme: Trypsin KR_C, number of missed cleavages: 3, precursor and fragment tolerance: +/-20ppm. The protein sequences of all identified proteins in pFind search, including indistinguishable proteins, were extracted from the original database to generate a pFind-generated reduced database using an in-house Perl script. For MSfragger search, FragPipe (v20.0) was used with either the full combined database or the pFind-generated reduced database for generating spectral libraries that will be used for analysis for DIA-PASEF dataset. For MSfragger search using the full combined database, a database split factor of 10 was used. Both searches followed the default workflow and using ciRT for spectral library generation. In addition to spectral libraries generated, the protein sequences of all identified proteins, including indistinguishable proteins, of MSFragger search against the original database were extracted using an in-house Perl script to generate a MSFragger derived reduced FASTA database.

#### DIA-NN search for identification and quantitation of DIA-PASEF data

DIA-PASEF data was processed with DIA-NN ^38^ (v1.8.1) using either spectral libraries generated through MSFragger or with a library-free mode using reduced FASTA databases generated through pFind or MSFragger searches as described above. Default settings were used for all the DIA-NN searches.

#### DDA-PASEF data quantitative search

To obtain identification and quantitation results of DDA-PASEF dataset, the DDA data was processed with the FragPipe (v21.1) MSFragger node using the LFQ-MBR workflow. The full mouse gut microbial gene catalog and host database was used. Similar to the spectral library generation workflow, a database split factor of 10 was used. MaxLFQ intensity of identified protein groups was used in this study with a minimum ion of 1.

#### Taxonomic and functional annotation and analysis

Taxonomic and functional annotation and analysis for both DIA-NN and MSFragger outputs was performed using MetaLab ^39^ (version 2.3). Briefly, the DIA-NN outputs report_pg.tsv and report_pr.tsv files were used for functional and taxonomic annotations, respectively. Similarly, the MSFragger outputs combined_proteins.tsv and combined_peptides.tsv were used for functional and taxonomic annotations, respectively. Unipept option in MetaLab was used to generate taxonomic annotations for both database search results, and a minimum of distinct peptide of 3 was used for confidently identification of taxa.

#### Database search for small proteins and atimicrobial peptides

A small protein database derived from human gut microbiome was downloaded from the supplemental data of a previous study by Sberro et. al. ^23^. We used the protein cluster data table which contained 444,054 entries to generate the small protein database. The AMPsphere AMP database was downloaded 2024/01/09 from AMPsphere (https://ampsphere.big-data-biology.org/home), containing 863,498 entries ^31^. To enable calculation of relative abundance of small protein or AMPs to total proteins in a sample, all identified peptide sequences from previous search using the combined database (gut microbial gene catalog and mouse proteome) were concatenated with the AMP or small protein databases for DIA-NN search or MSFragger search for DIA-PASEF data or DDA-PASEF data, respectively. DDA and DIA data was quantitatively analyzed using the methods described above with the respective alternate database.

### Data visualization and statistical analysis

Experimental flowcharts were generated using BioRender (https://www.biorender.com/). Boxplots, histograms, and pie charts were generated using R package *ggplot2* and *ggpubr*.

## DECLARATIONS

## Acknowledgments

We gratefully acknowledge Drs. Simon Sauvé, Huixin Lu and Michael Rosu-Myles from Health Canada for their critical comments on the manuscript.

## Authors’ contributions

Made substantial contributions to conception and design of the study and performed data analysis and interpretation: Wang A, Fekete E EF, Zhang X.

Performed data acquisition and contributed to data analysis and interpretation: Wang A, Fekete E EF, Zhang X, Creskey M, Cheng K, Ning Z, Figeys D.

Provided administrative, technical, and material support: Pfeifle A, Li X, Zhang X.

## Availability of data and materials

All MS proteomics data that support the findings of this study have been deposited to the ProteomeXchange Consortium (http://www.proteomexchange.org).

## Ethical approval and consent to participate

The animal procedures were approved by the Animal Care Committee in Health Canada and performed in accordance with institutional guidelines.

## Consent for publication

Not applicable.

