## SUPPLEMENTARY DATA for "Assessing fecal metaproteomics workflow and small protein recovery using DDA and DIA PASEF mass spectrometry"

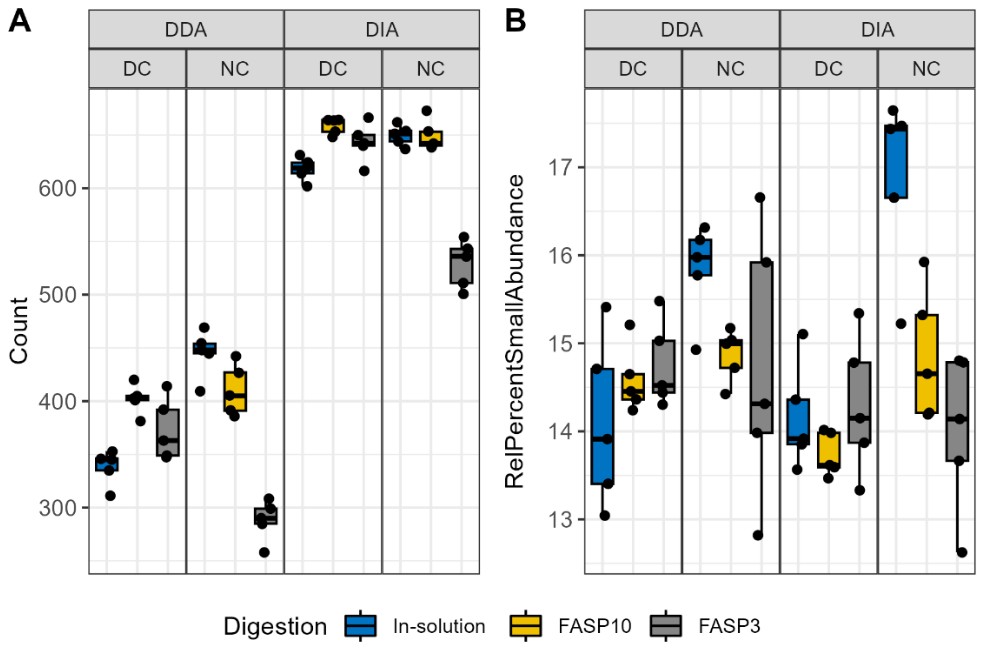


**Supplementary Figure 1.** Identification and abundance of host proteins in mouse fecal samples. Boxplots showing the number (A) of host proteins identified and their relative abundance (B) within the sample.

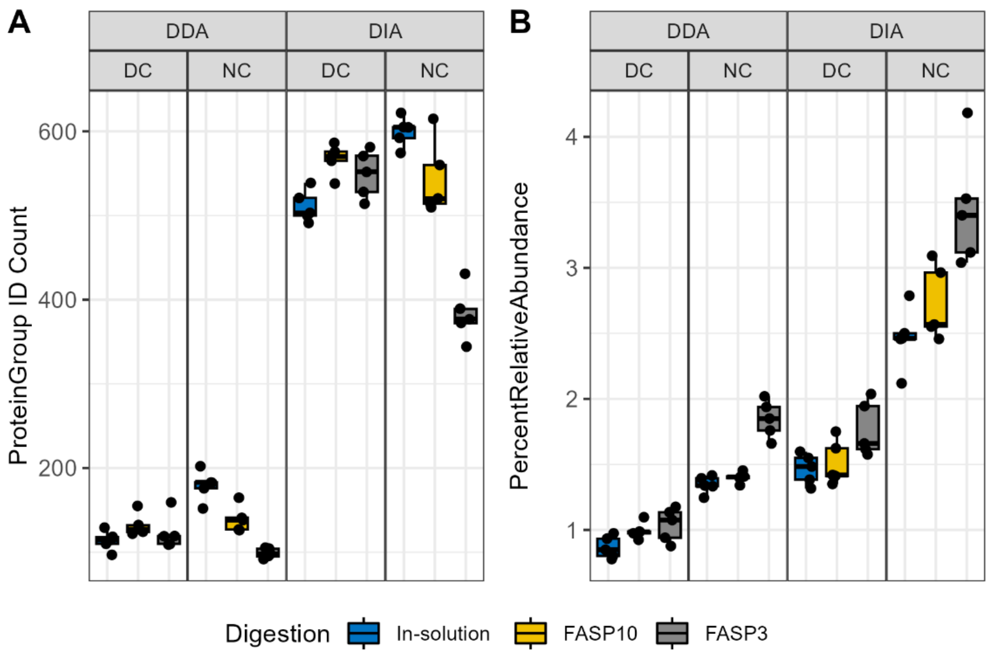


**Supplementary Figure 2.** Identification of small proteins in samples when searched against a small protein database. Count of identified small proteins (A). Percent relative abundance of small proteins compared to total abundance of proteins in sample (B).

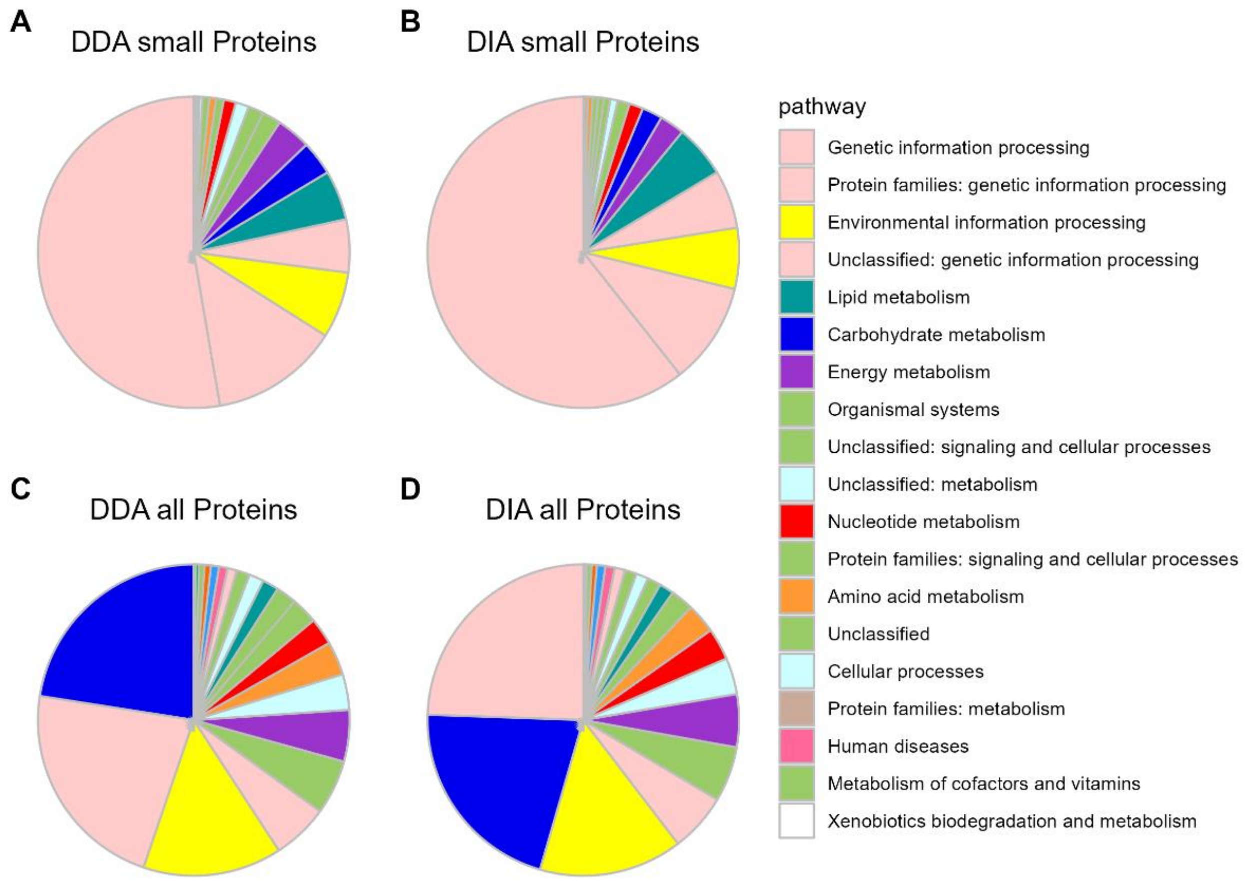


**Supplementary Figure 3**. GhostKOALA functional annotation of small proteins identified (≤ 100 amino acids) (A - B) as well as all proteins identified (C-D) in DDA and DIA datasets.

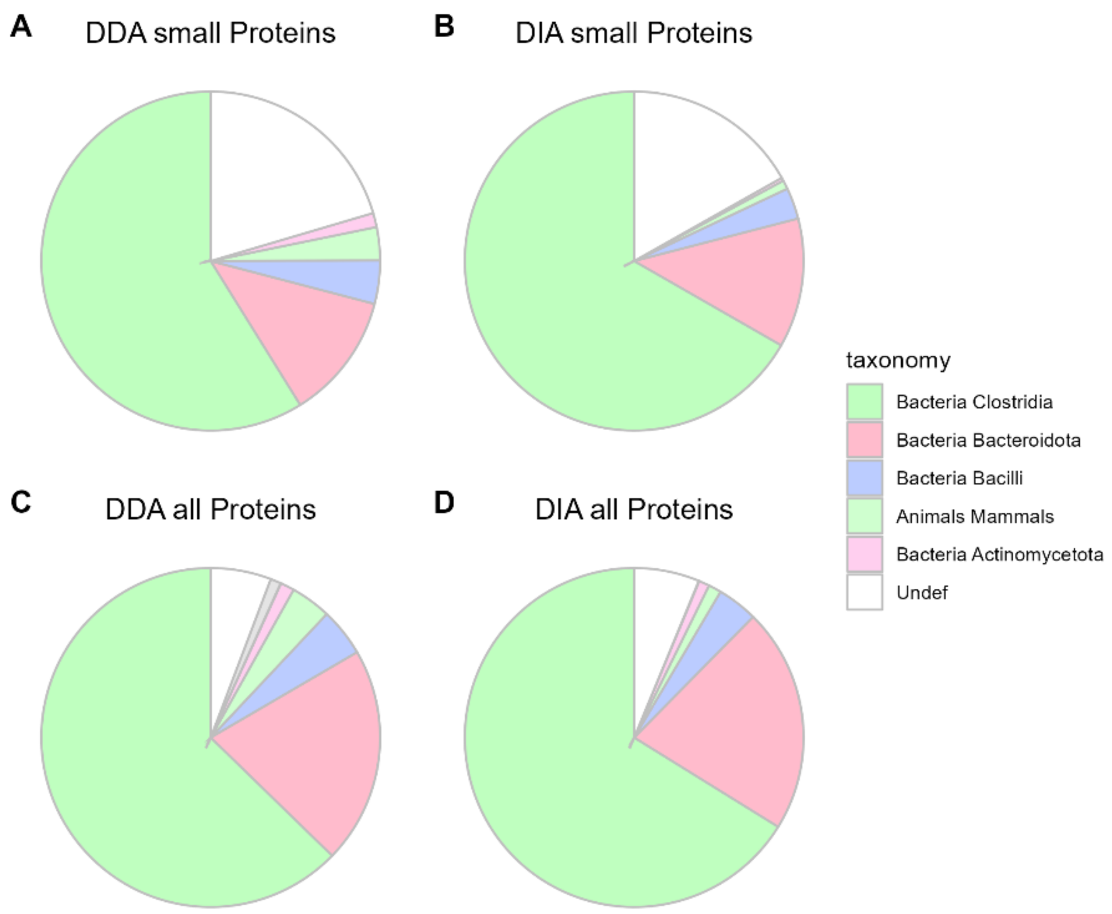


**Supplementary Figure 4**. GhostKOALA taxonomic annotation of small proteins identified (≤ 100 amino acids) (A - B) as well as all proteins identified (C-D) in DDA and DIA datasets.
